# A catalog of CasX genome editing sites in common model organisms

**DOI:** 10.1101/585521

**Authors:** Elisha D.O. Roberson

**Affiliations:** Department of Medicine, Division of Rheumatology, Washington University, St. Louis, MO 63110.; Department of Genetics, Washington University, St. Louis, MO 63110.

**Keywords:** genome editing, CasX, DpbCasX, Cas12e, *Saccharomyces cerevisiae*, *Caenorhabditis elegans*, *Drosophila melanogaster*, *Danio rerio*, *Mus musculus*, *Rattus norvegicus*, *Homo sapiens*

## Abstract

DpbCasX, also called Cas12e, is an RNA-guided DNA endonuclease isolated from *Deltaproteobacteria*. In this paper I characterized the CasX-compatible genome editing sites in the reference genomes of yeast (*Saccharomyces cerevisiae)*, flatworms (*Caenorhabditis elegans)*, flies (*Drosophila melanogaster*), zebrafish (*Danio rerio*), mouse (*Mus musculus*), rats (*Rattus norvegicus*), and humans (*Homo sapiens*). Across those genomes there were >27,000 CasX sites per megabase on average. More than 90% of genes in each genome had at least one unique site overlapping an exon, with median unique sites per gene of 6 – 45. I also annotated sites in the GRCm38 reference and 15 additional mouse strain genomes. The presence of specific guide sequences varied amongst the strains, with CAST/EiJ and PWK/PhJ showing the greatest divergence from the reference strain. The high density of CasX sites and number of exon overlapping sites suggests that CasX has the potential to be used as a common genome editor.

## 1. Introduction

Genome editing is a powerful molecular tool. The most widely used of these systems is the *Streptococcus pyogenes* clustered regularly interspaced short palindromic repeat (**CRISPR**)/Cas9 endonuclease (Jinek et al. 2012; Cong et al. 2013). Cas9 is an RNA-guided DNA endonuclease. One common goal of genome editing is to knockout a gene. A guide RNA targets the endonuclease complex to a specific genomic location, triggering a double-strand break. If the introduced breaks are repaired by non-homologous end joining (**NHEJ**) the result is a deletion of variable size. Similar techniques can be used to knock-in a specific genetic variant by simultaneously causing DNA damage while including a DNA oligonucleotide donor with the variant of interest (non-allelic homologous recombination; **NAHR**), or to cause transcriptional activation using a nuclease-dead Cas9 with a fused transcriptional activator (Maeder et al. 2013; Perez-Pinera et al. 2013; Wang et al. 2013). The knock-in method in particular has been touted as a potential mechanism to cure Mendelian-like genetic disorders by repairing the defective allele. A major concern for human gene-editing, however, is the possibility of introducing new genetic lesions through off-target activity of the editing enzyme. This has led to a general consensus that germline correction of a disease-causing variant in humans via Cas9 knock-in is not advisable at this time This has led to intense research into ways to reduce off-target editing for clinical use, and to identify smaller endonucleases to increase delivery efficiency.

These RNA-guided endonucleases require a protospacer adjacent motif (**PAM**) next to the DNA target sequence. Identical sequence matches without adjacent PAM sites are not efficiently cut. The PAM site for Cas9 is NGG, where N is any DNA base. One mechanism of reducing off-target effects would be to use an endonuclease with a longer PAM sequence. One recently identified RNA-guided genome editing endonuclease from *Deltaproteobacteria* is CasX (DpbCasX), tentatively designated as Cas12e (Koonin et al. 2017; Liu et al. 2019). This endonuclease has a 4 basepair PAM site (TTCN) with a 20 basepair guide sequence. It introduces a staggered double-strand break by cutting within the targeted sequence on the non-target strand and downstream of the target sequence on the target strand. The longer PAM site and smaller overall peptide size compared to Cas9 make CasX an attractive area of future genome editing development.

In this paper I present a catalog of CasX-compatible genome editing sites in 7 model organism genomes and in multiple mouse strains. The annotations are freely available from FigShare. Each site is annotated for chromosome, start and end position, PAM site, guide RNA target, uniqueness among editing sites, and any overlaps with the exons of known genes.

## 2. Materials and Methods

### 2.1. Identification of CasX sites

I coordinated part of the analysis using GNU Make (v4.1) on server running Ubuntu Linux (v16.04). The Makefile downloaded the gene annotations (GTF format) and genome sequences (FASTA format) for each of the specified genomes from release 95 of Ensembl (Flicek et al. 2011). The primary assembly FASTA file was used when it was available, and if none was found it fell back on using the top level file. I used the soft masked version (simple repeats as lower case characters rather than Ns) of the genome in either case. I calculated the GC content of each genome using a Python script with the pyfaidx package (v0.5.5.2) (Shirley et al. 2015). I then identified the potential CasX editing sites in each genome using Motif Scraper (v1.0.2) with the motif TTCNNNNNNNNNNNNNNNNNNNNN and multiple cores in file buffered mode (Roberson 2018). The standard Motif Scraper options buffer all sites in memory for sorting later based on the order of contigs in the input FASTA file. For large genomes this uses a considerable amount of RAM. In file buffered mode the hits are printed by contig and strand into temporary files that are concatenated at the end of the analysis to form the full output, substantially reducing the overall memory burden.

### 2.2. Site annotation

I performed the annotations on the Washington University Center for High Performance Computing cluster. The motif location output included the location of the hit in the genome (contig, start, end) and the associated sequences. I wanted to further analyze each output for uniqueness of the target site, PAM site usage, and overlap with the exons of known genes using R (v3.5.1) with the GenomicRanges, GenomicFeatures, here, knitr, multidplyr, and tidyverse packages (Lawrence et al. 2013). I calculated the size of the reference genomes from their FASTA index files, determined the uniqueness of guide RNA sites amongst all editing sites, counted PAM usage, and annotated any overlaps with the exons of known genes. It is worth noting that uniqueness of a guide was determined only among editing sites. Presumably another site that matches the guide exactly, but does not have the PAM site would not be cleaved. A guide that overlaps an exon can likely be used to knockout the gene or knock-in coding variants at the exon. Identifying unique editing sites that overlap promoters would require additional analysis. For the mouse strains, I calculated the presence (1) / absence (0) of guide sequences in each genome. I used this binary table to calculate the Hamming distance (Hamming 1950) between all strains in Python using pandas and the scipy spatial packages.

## 3. Results

### 3.1. CasX-compatible unique editing targets are common in model organism genomes

I was able to catalog and annotate potential CasX editing sites in 7 common model species: yeast (*Saccharomyces cerevisiae*), flatworms (*Caenorhabditis elegans*), fruit flies (*Drosophila melanogaster*), zebrafish (*Danio rerio*), mouse (*Mus musculus*), rats (*Rattus norvegicus*), and humans (*Homo sapiens*). I identified 263,193,973 total sites, of which 200,558,172 were unique cutters in their respective genomes (Table 1). Across the seven genomes there was an average of 1 site per 37 basepairs, and 1 unique site every 45 basepairs. The exon overlaps support the potential use of CasX to target genes of interest for editing. The median number of exon overlapping cutters per gene ranged from 7 to 52 across the genomes tested, with between 6 and 45 unique cutters per gene (Table 2). Importantly, at least 90% of annotated genes across all organisms had at least one unique CasX site overlapping at least one exon. There were more A/T PAM sites (TTCA, TTCT) compared to C/G (TTCC, TTCG) PAM sites in all the surveyed genomes (Fig. 1). The TTCG PAM site is particularly depleted in zebrafish, mouse, rat, and human genomes, perhaps due to the general depletion of CpG sites genome-wide.

**Table 1.** CasX site genomic distribution in 7 model organisms

| Organism | Total sites | Unique sites | Unique (%) | Total sites / Mbp<br>[median] | Unique sites / Mbp<br>[median] |
| --- | --- | --- | --- | --- | --- |
| *S. cerevisiae* | 367,810 | 345,376 | 93.90 | 30,254.74 | 28,409.4 |
| *C. elegans* | 3,315,259 | 2,988,557 | 90.15 | 33,057.91 | 29,800.2 |
| *D. melanogaster* | 3,681,483 | 3,011,422 | 81.80 | 25,614.59 | 20,952.5 |
| *D. rerio* | 33,296,705 | 22,500,532 | 67.58 | 24,242.74 | 16,382.2 |
| *M. musculus* | 70,817,235 | 54,464,834 | 76.91 | 25,932.10 | 19,944.1 |
| *R. norvegicus* | 73,427,248 | 55,157,865 | 75.12 | 25,582.77 | 19,217.5 |
| *H. sapiens* | 78,288,233 | 62,089,586 | 79.31 | 25,256.30 | 20,030.5 |

**Table 2.** CasX sites overlapping known gene exons

| **Organisms** | **Genes** | **Cut (%)** | **Unique cut (%)** | **Sites / gene** | **Unique sites / gene** |
| --- | --- | --- | --- | --- | --- |
| *S. cerevisiae* | 7,036 | 99.97 | 96.93 | 31 | 29 |
| *C. elegans* | 46,778 | 96.46 | 94.73 | 7 | 6 |
| *D. melanogaster* | 17,737 | 99.86 | 97.30 | 38 | 37 |
| *D. rerio* | 32,520 | 99.94 | 95.43 | 52 | 45 |
| *M. musculus* | 54,838 | 99.58 | 93.14 | 36 | 26 |
| *R. norvegicus* | 32,883 | 99.40 | 90.44 | 33 | 25 |
| *H. sapiens* | 58,735 | 99.60 | 96.98 | 28 | 22 |

**Fig. 1.**
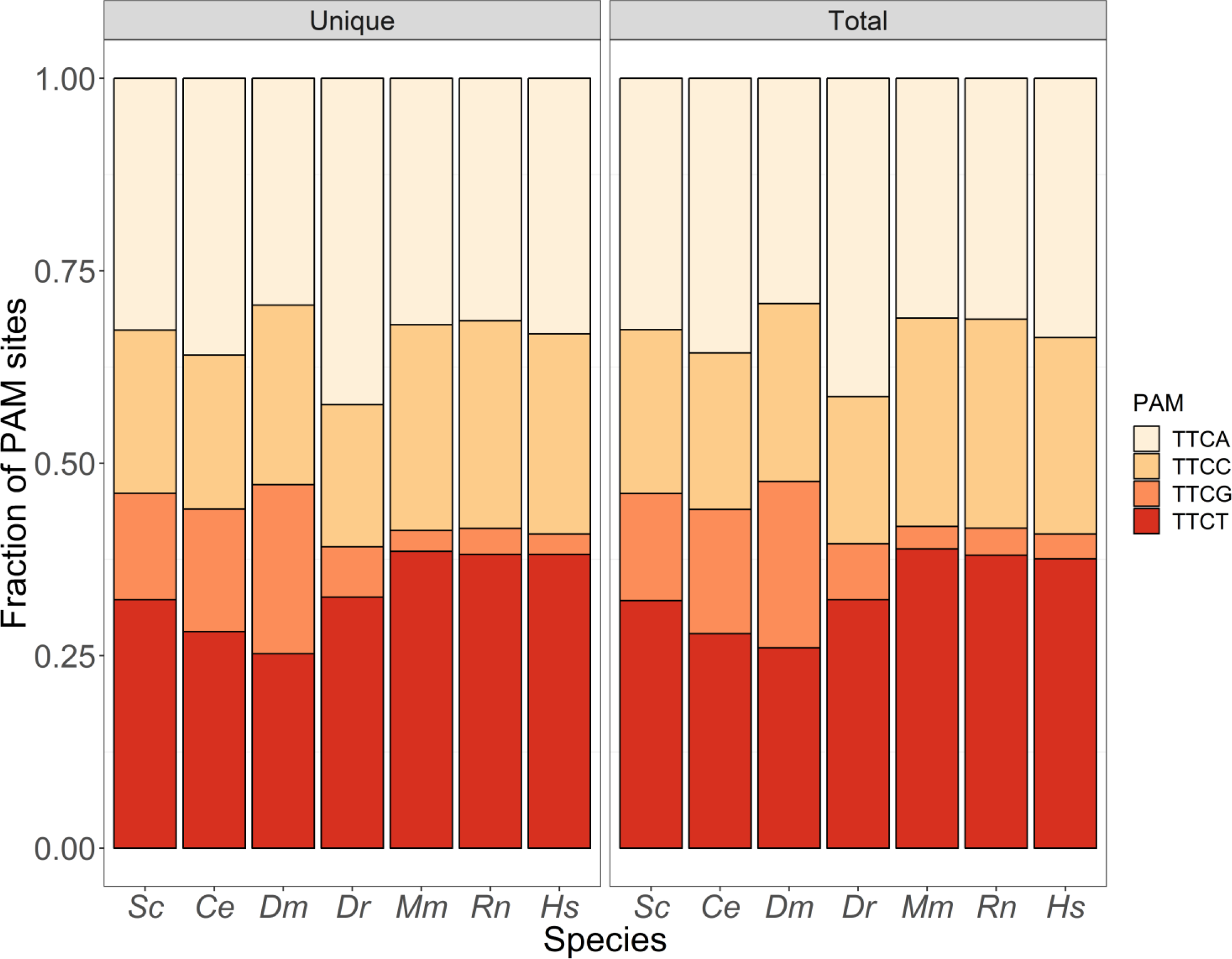
CasX PAM site usage. Shown in this figure are the 7 species on the x-axis (abbreviated as the first letter of the genus and species), and a stacked bar chart of fractional PAM site usage on the y-axis. The plot is divided into two subplots with the distribution of only unique cutters and of all sites. The A and T PAM sites are generally the most used. The TTCC and TTCG sites are used much less often in zebrafish, mouse, rat, and humans. The TTCG site, which contains a CpG dinucleotide, is seldom observed in those four species in particular.

### 3.2. Mouse strains vary in guide RNA site availability in their reference genomes

I also annotated the CasX sites in the main Ensembl mouse reference (GRCm38; *Mus musculus*) and in multiple strains: 129S1/SvImJ, A/J, AKR/J, BALB/cJ, C3H/HeJ, C57BL/6NJ, CAST/EiJ, CBA/J, DBA/2J, FVB/NJ, LP/J, NOD/ShiLtJ, NZO/HlLtJ, PWK/PhJ, and WSB/EiJ. The GRCm38 reference is built primarily from the C57BL/6J strain. Genome-editing is especially important in the generation of mouse models, drastically reduced the time and effort required to generate knockouts and knock-ins. However, databases of editing sites for mice are built mostly from the GRCm38 reference, and therefore most applicable to C57BL/6J.

I used a binary table of presence / absence of guide RNA sequences across all 16 genomes to calculate the Hamming distance between all pair-wise strain comparisons (Fig. 2). One important finding is that there is different site availability in the strains. The GRCm38 reference and C57BL/6NJ had few differences (distance 0.017), as expected. The site availability was also similar between 129S1/SvImJ and LP/J (distance 0.038). Two strains stood out as most different from the C57BL/6J reference, CAST/EiJ (distance 0.297) and PWK/PhJ (distance 0.291), and from each other (distance 0.289). These strains in particular might benefit from strain-specific guide RNA development.

**Fig. 2.**
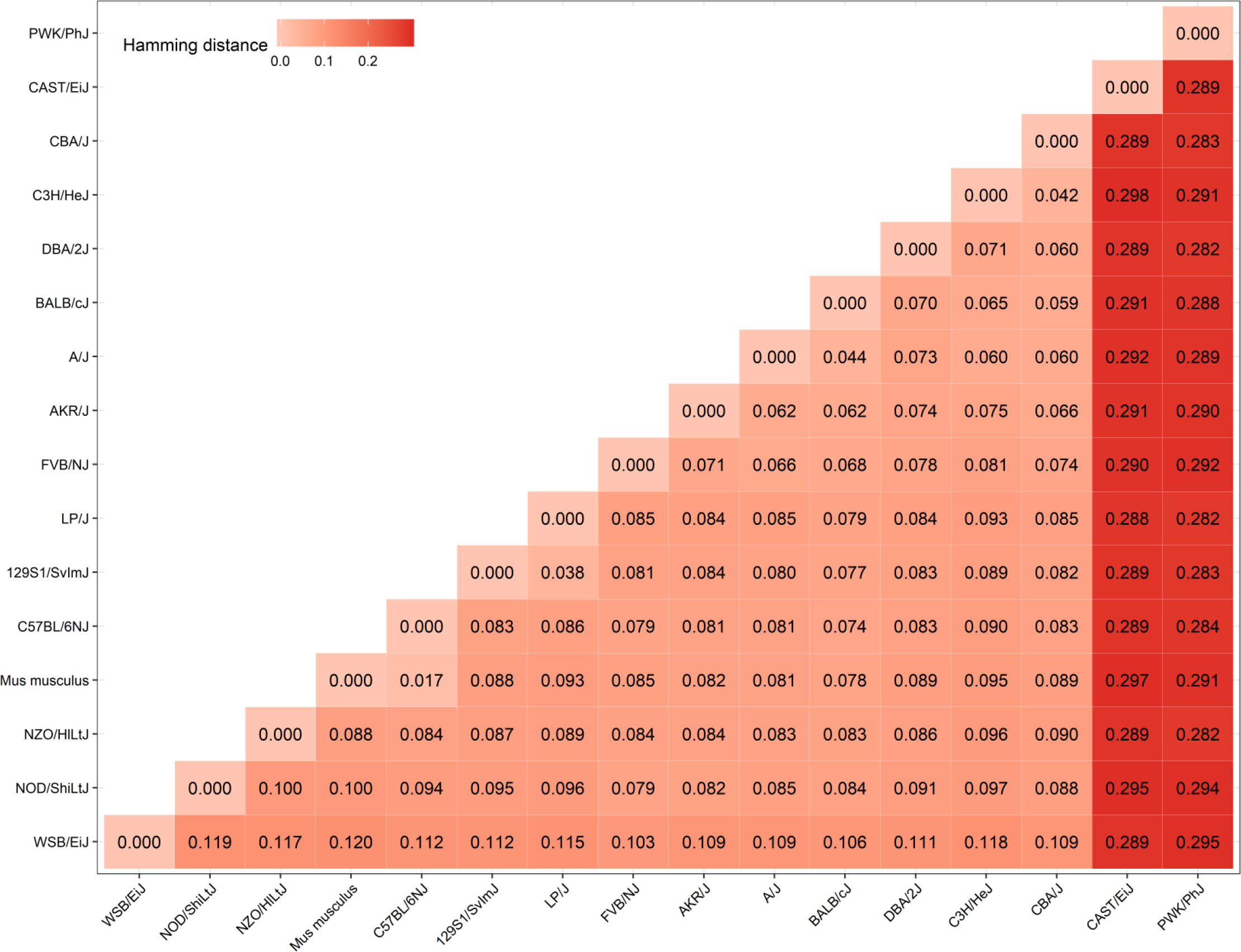
Hamming distance for editing targets amongst Ensembl mouse strains. Shown is the Hamming distance between sites for the GRCm38 reference (denoted *Mus musculus*), and genomes of 15 strains available via Ensembl. The matrix was ordered by calculating the hierarchical clustering of the distances with complete linkage. The strains CAST/EiJ and PWK/PhJ were the furthest from the GRCm38 reference among the tested strains, though all had some degree of difference. Those two strains in particular might require site annotations for their specific genomes rather than site selection from the general mouse reference.

## 4. Discussion

Identifying new RNA-guided endonucleases to use as genome editors is an area of intense research. There are currently many modifications of Cas9 that can help to decrease the number of off-target cuts (such as Cas9 nickase), but it is still favorable to explore other editors with more favorable characteristics for clinical use. CasX appears to use a mechanism distinct from both Cas9 and Cas12a, suggesting it may have different benefits and limitations (Liu et al. 2019). CasX guide sites are relatively common in all the tested genomes, and most genes have at least one CasX site overlapping an exon. This supports the potential utility of CasX in genome editing. The expanded PAM site also may reduce the number of off-target near matches in candidate genomes. Amongst the mouse strains, there were some substantial differences in site availability. In some strains, particularly CAST/EiJ and PWK/PhJ, there are many differences in site availability between them and the GRCm38 reference. It is important to note that the resolution of these differences is directly dependent on the quality of genome assembly. Any strains with poor assembly may have dropout of sites that is technical rather than biological. Regardless, this catalog of CasX editing sites will be an important resource in the future testing of this new class of RNA-guided genome editor.

## 5. Data availability

The code to generate this analysis is available on GitHub: https://github.com/RobersonLab/2019CasXModelOrgCatalog

The cataloged CasX sites are available on FigShare collected as a project: https://figshare.com/projects/2019_CasX_genome_editing_site_annotations/61103

## 6. Acknowledgements

E.D.O.R. was partially supported by NIH grant P30-AR073752. Some of the computations in this paper were performed using the facilities of the Washington University Center for High Performance Computing, which were partially provided through NIH grant S10 OD018091.

## 10. Supplemental tables

**Supp. Table 1.**
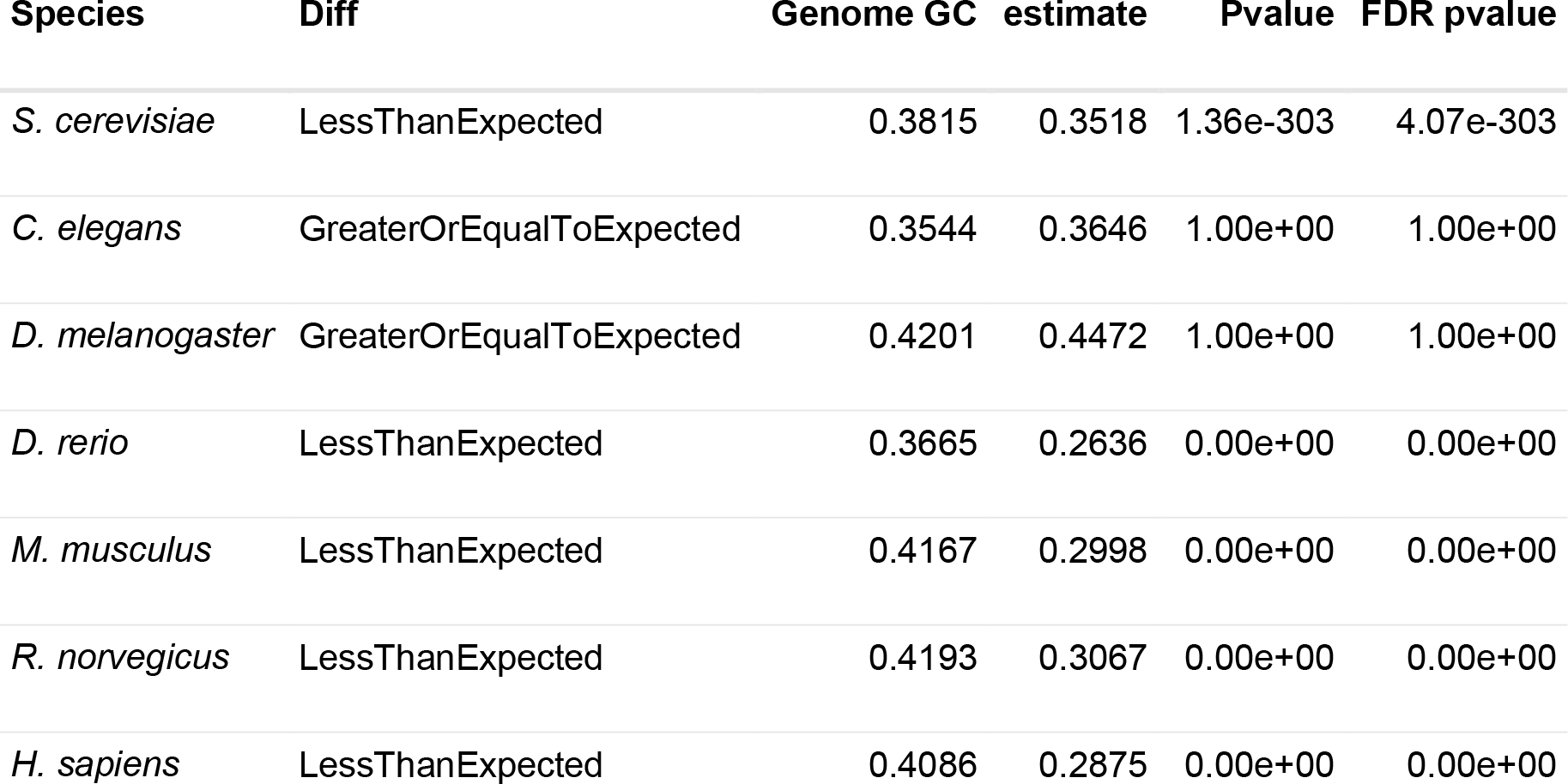
GC PAM sites depleted in reference genomes. Genome GC was calculated based on the genome FASTA file. The estimate is the binomial test estimate of actual C/G usage in the PAM sites, with the associated p-value and the Benjamini Hochberg FDR p-value.

